# Variance and limiting distribution of coalescence times in a diploid model of a consanguineous population

**DOI:** 10.1101/2020.06.30.180521

**Authors:** Alissa L. Severson, Shai Carmi, Noah A. Rosenberg

## Abstract

Recent modeling studies interested in runs of homozygosity (ROH) and identity by descent (IBD) have sought to connect these properties of genomic sharing to pairwise coalescence times. Here, we examine a variety of features of pairwise coalescence times in models that consider consanguinity. In particular, we extend a recent diploid analysis of mean coalescence times for lineage pairs within and between individuals in a consanguineous population to derive the variance of coalescence times, studying its dependence on the frequency of consanguinity and the kinship coefficient of consanguineous relationships. We also introduce a separation-of-time-scales approach that treats consanguinity models analogously to mathematically similar phenomena such as partial selfing, using this approach to obtain coalescence-time distributions. This approach shows that the consanguinity model behaves similarly to a standard coalescent, scaling population size by a factor 1 − 3*c*, where *c* represents the kinship coefficient of a randomly chosen mating pair. It provides the explanation for an earlier result describing mean coalescence time in the consanguinity model in terms of *c*. The results extend the potential to make predictions about ROH and IBD in relation to demographic parameters of diploid populations.

## 1 Introduction

Previously (Severson *et al.*, 2019), we devised a coalescent model of a consanguineous diploid population in order to jointly study runs of homozygosity (ROH) and identity-by-descent (IBD) sharing. We used the fact that the time to the most recent common ancestor at a locus for a pair of genomes is inversely related to the length of the shared segment around the locus (Palamara *et al.*, 2012; Carmi *et al.*, 2014). To distinguish within-individual from between-individual pairwise coalescence times, which describe levels of ROH and IBD sharing, respectively, we modeled a diploid biparental population.

We developed our model by extending the diploid sib mating model of Campbell (2015) to allow *n*th cousin mating and a superposition of multiple degrees of cousin mating. To derive mean pairwise coalescence times in our models, we performed a first-step analysis of a Markov chain to condition on the state of two alleles in prior generations. We found that owing to the possibility of extremely recent coalescence from consanguinity, the effect of consanguinity reduced mean pairwise coalescence times within an individual as well as between separate individuals. The reduction of the mean coalescence time was proportional to the kinship coefficient of a randomly chosen mating pair, with a greater reduction for two alleles within an individual versus between individuals.

Although mean coalescence times are useful for summarizing the effect of consanguinity under the model, and they reveal that both within- and between-individual coalescence times depend on population size and consanguinity, the mean describes only one aspect of the distribution. Distributions of pairwise coalescence times within and between individuals are needed to more fully understand the effect of consanguinity on the length of shared segments within and between individuals (Palamara *et al.*, 2012; Carmi *et al.*, 2014).

Coalescent-based population-genetic models typically approximate a two-sex diploid population of size *N* individuals with a haploid population of size 2*N*. In a diploid model, two alleles can be either within an individual or in separate individuals, whereas in a haploid model, all alleles are exchangeable. Despite this distinction, with a sufficiently large population size and equal numbers of males and females, the ancestral process of a diploid population converges to that of a haploid population of twice the size, supporting the approximation (Wakeley, 2009, Chapter 6.1). Intuitively, the convergence occurs when the probability of rapid coalescence of the two alleles in a diploid individual is negligible.

A variety of techniques have been used to analyze coalescent models, in which, like in consanguinity models, rapid coalescence cannot be ignored. Wakeley *et al.* (2012) examined the effect of a fixed diploid population pedigree on coalescence times, finding that the probability of recent coalescence events is slightly increased compared to a corresponding haploid model. Models of partial selfing (Nordborg and Donnelly, 1997; MÖhle, 1998b) have examined the effect of a selfing rate that gives the probability of immediate coalescence of pairs of alleles in a diploid individual. This work has used the separation-of-time-scales approach (MÖhle, 1998b), which describes the concurrent effect of a “fast” coalescent process (coalescence of two alleles from an individual due to selfing) and a “slow” process (coalescence of pairs of alleles in the population at large).

Here, we continue our earlier work to further interrogate the distributions of coalescence times for pairs of alleles within and between individuals in a diploid coalescent model with consanguinity. First, we derive the variance of coalescence times under sib mating and extend the calculation to the superposition case. Next, we use a separation-of-time-scales approach to derive the distributions of pairwise coalescence times within and between individuals in the limit of large population size. We compare the mean and variance of these limiting distributions to the exact solutions. We also compare the full limiting distribution to numerical solutions in the sib mating case, and also to the results of simulations from the exact Markov chains. We find that the limiting distributions closely approximate the exact distributions.

## 2 Model

We extend the model of Severson *et al.* (2019), which itself extended the model of Campbell (2015). The model considers a constant-sized diploid population with discrete generations. Each generation has *N* ≥ 2 monogamous mating pairs, 2*N* diploid individuals, and 4*N* alleles at each locus. A constant fraction of the mating pairs are consanguineous unions, and the other pairs are non-consanguineous. The fraction of mating pairs each generation that are related as *n*th cousins is denoted *c*_*n*_ (Figure 1A).

**Figure 1:**
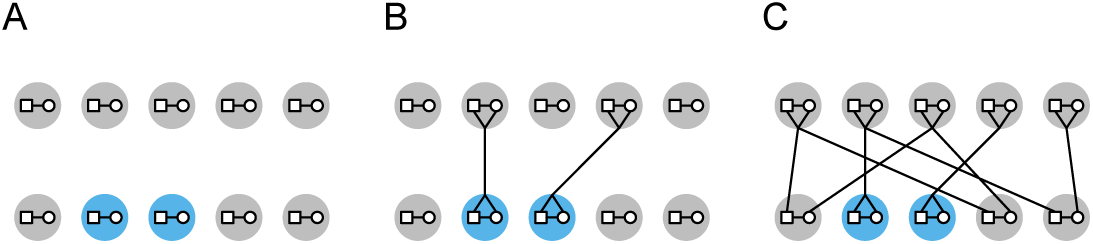
Diploid model of sib mating. **(A)** In each generation, a fraction *c*_0_ = 0.4 of *N* = 5 mating pairs are sib mating pairs. **(B)** Sib mating pairs are each assigned a parental pair from the previous generation. **(C)** Non-consanguineous pairs are each assigned two distinct parental pairs, representing the two sets of parents for the two individuals in the non-consanguineous pair.

To illustrate the model, we consider a simple case, a population with sib mating, viewing sibs as 0th cousins. Backward in time, the *c*_0_*N* sib mating pairs—each of whose two individuals necessarily share a single set of parents—each randomly choose one parental mating pair from the previous generation (Figure 1B). Next, the (1 − *c*_0_)*N* random-mating pairs each randomly choose two distinct parental mating pairs (Figure 1C). Note that two individuals in a random-mating pair cannot share the same parental mating pair, so that chance sib mating is forbidden. Because parental mating pairs are chosen at random, two individuals in separate mating pairs will be siblings with probability 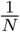.

In this model, two alleles at a locus can be in three possible states, denoted 1, 2, and 3. State 1 corresponds to two alleles within an individual, state 2 is for two alleles in two distinct individuals in a mating pair, and state 3 is two alleles in two distinct individuals in separate mating pairs. We use the random variables *T, U*, and *V* to denote the coalescence times for two alleles in states 1, 2, and 3, respectively.

## 3 Variance of coalescence times

### 3.1 Sib mating

We begin with a sib mating population. In each generation, a constant fraction *c*_0_ of the *N* mating pairs are siblings, and chance sib mating is forbidden. Previously, we derived the means for *T, U*, and *V* as (Severson *et al.*, 2019, eqs. 4-6)

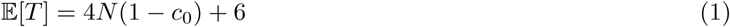

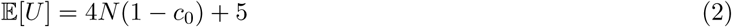

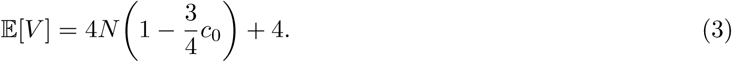

Here we derive the variances for *T, U*, and *V*. First, two alleles that within an individual (state 1) are always in two individuals in a mating pair in the previous generation (state 2), so *T* = *U* + 1 and

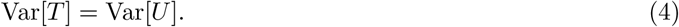

Next we derive Var[*U*] using the law of total variance. For convenience, we define the random variable *Z* to be the state of two alleles in the previous generation. We add state 0 to represent coalescence, so *Z* takes values from {0, 1, 2, 3}. Applying the law of total variance and conditioning on *Z*, Var[*U*] = 𝔼[Var[*U* |*Z*]] + Var[𝔼[*U* |*Z*]].

Beginning with 𝔼[Var[*U* |*Z*]], if two alleles are in a mating pair (state 2), then the previous generation has four possible states, encoded in values of *Z*. First, with probability *c*_0_, the mating pair is a sib mating pair. In this case, with probability 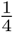, the alleles coalesce in the previous generation (*Z* = 0) and the alleles have coalescence time 1. Similarly, if the mating pair is consanguineous, then with probability 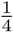, the alleles were inherited from the same individual (*Z* = 1), and they have coalescence time *T* + 1. Lastly if the mating pair is a sib mating pair, then with probability 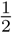, the two alleles were inherited from separate parents in the previous generation (*Z* = 2), and the two alleles have coalescence time *U* + 1. With probability 1 − *c*_0_, the two alleles are in a random-mating pair, in the previous generation they were in two individuals in separate mating pairs (*Z* = 3), and they have coalescence time *V* + 1. Combining these cases gives

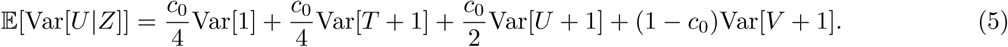

For the next term, Var[𝔼[*U* |*Z*]], we rewrite it using the definition of variance, Var[𝔼[*U* |*Z*]] = 𝔼[𝔼[*U* |*Z*]^2^] − 𝔼[𝔼[*U* |*Z*]]^2^ = 𝔼[𝔼[*U* |*Z*]^2^] − 𝔼[*U*]^2^. Because 𝔼[*U*]^2^ is known (eq. 2), we only need 𝔼[𝔼[*U* |*Z*]^2^], which we derive by again conditioning on *Z*. With probability *c*_0_*/*4, the alleles coalesce in 1 generation (*Z* = 0). With probability *c*_0_*/*4, the alleles were inherited from the same individual (*Z* = 1), and 𝔼[*U* |*Z* = 1]^2^ = 𝔼[*T* + 1]^2^. With probability *c*_0_*/*2, the alleles were inherited from separate individuals in the same mating pair (*Z* = 2), and 𝔼[*U* | *Z* = 2]^2^ = 𝔼[*U* + 1]^2^. Lastly, with probability 1 − *c*_0_, the alleles are in a non-consanguineous mating pair and they were inherited from two individuals in separate mating pairs (*Z* = 3), giving 𝔼[*U* | *Z* = 3]^2^ = 𝔼[*V* + 1]^2^. Combining these cases gives

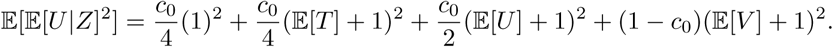

Subtracting 𝔼[*U*]^2^, we have

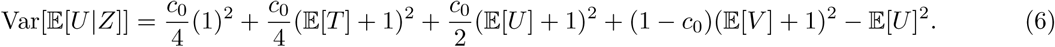

Summing eqs. 5 and 6, applying eq. 4 and 𝔼[*T*] = 𝔼[*U*] + 1, and simplifying gives

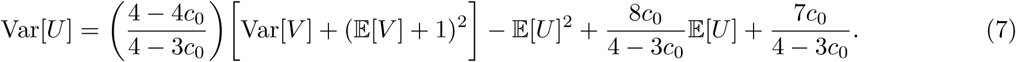

Next, for Var[V], we again use the law of total variance and condition on *Z*. For the first term 𝔼[Var[*V* | *Z*]], recall that if two alleles are in two separate mating pairs, then because parents are chosen randomly with replacement, the two individuals are siblings with probability 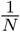. Then the probability that the two alleles are in two individuals who are siblings and that those alleles coalesce in the previous generation (*Z* = 0) is 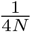. Similarly, the probability that the siblings inherit distinct alleles from the same parent (*Z* = 1) is 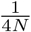, giving a coalescence time of *T* + 1. If the alleles are inherited from separate parents (*Z* = 2), an event with probability 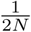, then the coalescence time is *U* + 1. Lastly, with probability 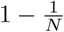, the individuals are not siblings, so the alleles were inherited from separate individuals in separate mating pairs (*Z* = 3), giving coalescence time *V* + 1. These four cases give

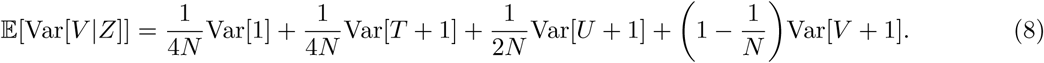

For Var[𝔼[*V* | *Z*]] = 𝔼[𝔼[*V* | *Z*]^2^] − 𝔼[𝔼[*V* | *Z*]]^2^, we have 𝔼[𝔼[*V* | *Z*]] = 𝔼[*V*] as before. For 𝔼[𝔼[*V* | *Z*]^2^], if two alleles are in two individuals in separate mating pairs, then with probability 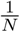, the individuals are siblings. If *Z* = 0, then the alleles coalesce in the previous generation with probability 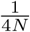. If *Z* = 1, then the two alleles were inherited from the same individual in the previous generation. This event occurs with probability 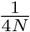, and 𝔼[*V* |*Z* = 1]^2^ = 𝔼[*T* + 1]^2^. With probability 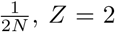, and the alleles were inherited from two individuals in a mating pair, giving 𝔼[*V* |*Z* = 2]^2^ = 𝔼[*U* + 1]^2^. Lastly, with probability 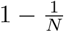, the two individuals are not siblings, so the alleles were inherited from two individuals in separate mating pairs (*Z* = 3), and 𝔼[*V* |*Z* = 3]^2^ = 𝔼[*V* + 1]^2^. Combining cases and subtracting 𝔼[*V*]^2^ gives the second term

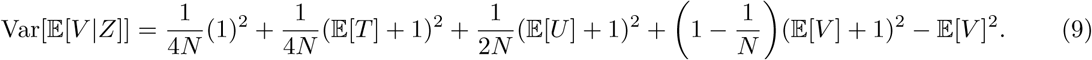

Summing eqs. 8 and 9, applying eq. 4 and 𝔼[*T*] = 𝔼[*U*] + 1, and simplifying gives the form

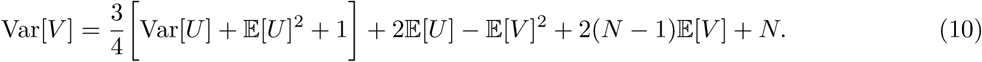

Eqs. 4, 7, and 10 form a linear system in Var[*T*], Var[*U*], and Var[*V*], which we solve, applying eqs. 1-3:

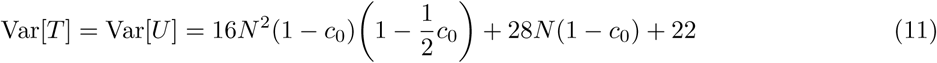

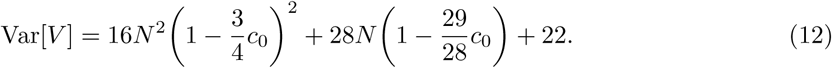

Eqs. 11 and 12 give the desired variances. We can immediately make a number of observations.

First, considering all possible consanguinity levels *c*_0_, both eqs. 11 and 12 are maximized when *c*_0_ = 0, and they decrease with increasing *c*_0_. Thus, consanguinity decreases variance for all three coalescence times.

Next, the difference Var[*V*] − Var[*T*] equals 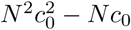, which is positive for 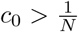, so that Var[*V*] > Var[*T*]. Thus, with a nontrivial consanguinity level, the variance of the coalescence time for two alleles in separate mating pairs exceeds that for two alleles in the same individual.

Third, taking *N* → ∞, eqs. 11 and 12 give

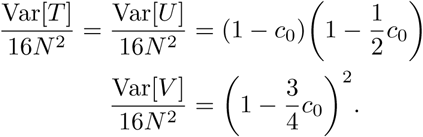

Thus, for sufficiently large *N*, the variances are dominated by the product of 16*N* ^2^, the variance of coalescence time for a haploid population of size 4*N*, and a reduction factor, (1 − *c*_0_) 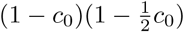 in eq. 11 and 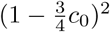 in eq. 12.

Using eqs. 3 and 12, note that in the limit of large *N*, Var[*V*]*/* 𝔼[*V*]^2^ → 1. Recall that an exponentially distributed random variable with mean *λ* has variance *λ*^2^, so the variance is the square of the mean. Although this relationship is not unique to the exponential distribution, the fact that Var[*V*]*/* 𝔼[*V*]^2^ → 1 is consistent with *V* being exponentially distributed in the limit as *N* → ∞. On the other hand, Var[*T*]*/* 𝔼[*T*]^2^ → (2 − *c*_0_)*/*(2 − 2*c*_0_), so *T* is not exponentially distributed in the *N* → ∞ limit for *c*_0_ > 0.

Eqs. 11 and 12 normalized by 16*N* ^2^ are plotted in Figure 2 as a function of the number of mating pairs *N* and fraction of sib mating pairs *c*_0_. As population size increases, the normalized variances quickly approach the reduction factors, 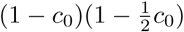 in eq. 11 and 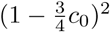 in eq. 12.

**Figure 2:**
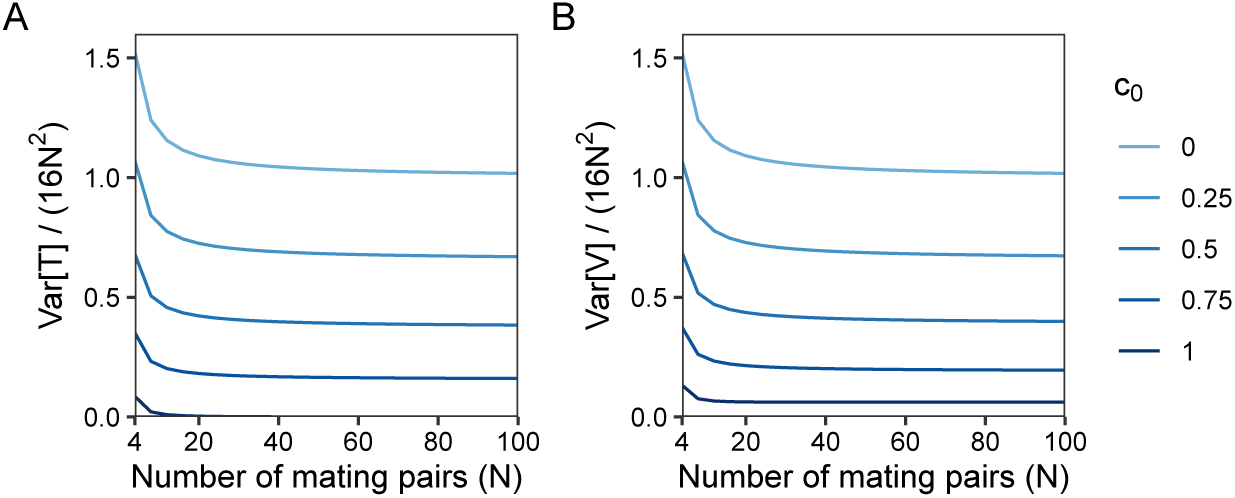
Normalized variance of pairwise coalescence times as a function of the number of mating pairs *N* and the fraction of sib mating pairs *c*_0_. **(A)** Var[*T*]*/*(16*N* ^2^), eq. 11. **(B)** Var[*V*]*/*(16*N* ^2^), eq. 12.

### 3.2 Superposition of multiple mating levels

In this section, we generalize the variance under sib mating to a superposition of multiple levels of cousin mating. Under the superposition, *i*th cousin mating is permitted for all *i* from 0 to *n*, where *n* is the degree of the most distant permissible cousin relationship. The case of *i* = 0 corresponds to sib mating. For each *i*, let *c*_*i*_ be the fraction of *i*th cousin mating pairs in each generation. For each *i* ≤ *n*, chance *i*th cousin mating is prohibited. We assume individuals in a consanguineous mating pair share only one line of descent—that is, for example, they cannot be both first and third cousins.

Under this model, we derived the means for *T, U*, and *V* as (Severson *et al.*, 2019, eqs. 17-19)

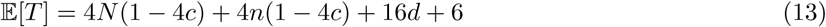

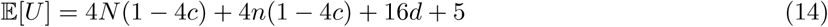

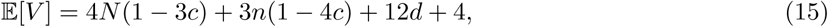

where *c*, the kinship coefficient for two individuals in a mating pair, is defined as

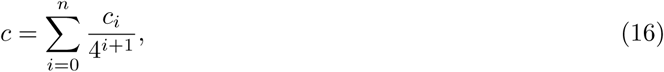

and for convenience, we define *d* as

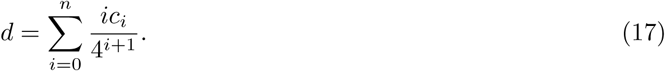

First for Var[*T*], as before, two alleles present within an individual are in two individuals in a mating pair in the previous generation, so *T* = *U* + 1 and eq. 4 continues to hold.

To derive Var[*U*], we again use the law of total variance and condition on *Z*. Before, if two individuals in a mating pair were siblings, then the alleles could transition to one of four states in the previous generation. Under the superposition, there are instead 3(*n* + 1) + 1 possible transitions. For each *i*, 0 ≤ *i* ≤ *n*, if the two individuals in the mating pair are *i*th cousins, then *i* + 1 generations in the past, the two alleles are inherited from the shared ancestral mating pair with probability 1*/*4^*i*^. If the alleles are inherited from the shared ancestral mating pair, then *i* + 1 generations ago they can transition to states 0, 1, or 2, giving 3(*n* + 1) possible transitions when considering all *i* from 0 to *n*. If the two individuals in the mating pair are not related, then because chance *n*th cousin mating is forbidden, *n* + 1 generations ago the alleles are in state 3, accounting for the last of the 3(*n* + 1) + 1 transitions.

For the first term in the law of total variance, 𝔼[Var[*U* | *Z*]], we consider *i*th cousin mating for each *i*. With probability *c*_*i*_*/*4^*i*^, the individuals are *i*th cousins and the two alleles were inherited from the shared ancestral mating pair *i* + 1 generations ago. If the alleles are inherited from this mating pair, then with probability 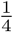, they coalesce in time *i* + 1; with probability 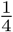, the alleles are inherited from the same individual in the mating pair, with coalescence time *T* + *i* + 1; and with probability 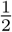, the alleles are inherited from separate individuals in the shared ancestral mating pair and have coalescence time *U* + *i* + 1. If for all *i*, 0 ≤ *i* ≤ *n*, the two individuals are not *i*th cousins, or if they are *i*th cousins for some *i* but the alleles are not inherited from the shared ancestral mating pair, then they are in two individuals in separate mating pairs *n* + 1 generations ago; this event has probability 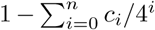, and the alleles have coalescence time *V* + *i* + 1. Summing the probabilities of the cases for 0 ≤ *i* ≤ *n*,

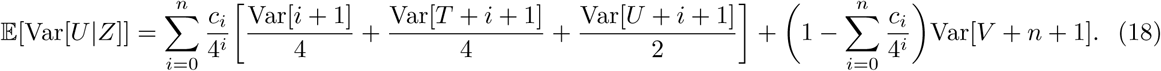

For the second term Var[𝔼[*U* | *Z*]], we derive 𝔼[𝔼[*U* | *Z*]^2^]. Again if the two individuals in the mating pair are *i*th cousins, then with probability *c*_*i*_*/*4^*i*^, the alleles were inherited from the shared ancestral mating pair *i*+1 generations ago. If the alleles were inherited from the ancestral mating pair, then there are three possible transitions: with probability 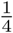, they coalesce with time *i* + 1; with probability 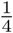, they were inherited from the same individual, giving mean 𝔼[*T* + *i* + 1]^2^; with probability 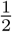, they were inherited from the two separate individuals, with mean 𝔼[*U* + *i* + 1]^2^. With probability 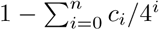, the alleles were not inherited from the shared ancestral mating pair for any *i*, 0 ≤ *i* ≤ *n*, and the alleles are not in a consanguineous mating pair, and then *n* + 1 generations ago they are in separate mating pairs, giving mean 𝔼[*V* + *n* + 1]^2^. Summing these cases over all *i* and subtracting 𝔼[*U*]^2^ (eq. 14) gives the second term

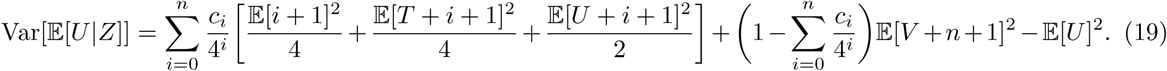

For Var[*V*], because parental mating pairs are chosen randomly with replacement, eq. 10 continues to hold. Hence, the sum of eqs. 18 and 19 gives Var[*U*], which together with eqs. 4 and 10 forms a linear system of equations. Applying eqs. 13-15, the solution is

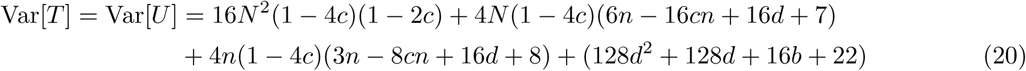

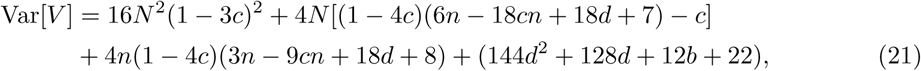

where

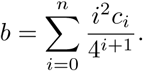

We can quickly observe that if *c* = *c*_*i*_*/*4^*i*+1^ for any *i*, then eqs. 20 and 21 reduce to the equations for Var[*T*] = Var[*U*] and Var[*V*] for *i*th cousin mating. In particular, if *c* = *c*_0_*/*4, then *d* = 0 and *b* = 0, and eqs. 20 and 21 reduce to eqs. 11 and 12, respectively.

Taking *N* → ∞, eqs. 20 and 21 give

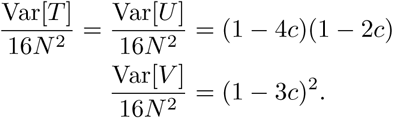

For large population size, the variances approach the product of 16*N* ^2^, the variance of coalescence time in a haploid population of size 4*N*, and reduction factors (1 − 4*c*)(1 − 2*c*) in eq. 20 and (1 − 3*c*)^2^ in eq. 21. For large *N*, Var[*V*] − Var[*T*] ≈ 16*N*^2^*c*^2^, a quantity that increases with consanguinity *c*.

Note that because 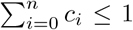, the maximum of *c* over all possible vectors (*c*_0_, *c*_1_, *…, c*_*n*_) is found by setting *c*_0_ = 1. The maxima for *d* and *b* set *c*_1_ = 1, because the *i* = 0 terms are 0 for *d* and *b*. Then 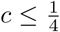, 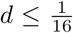, and 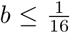. Assuming *n* ≪*N*, terms with constants *c, d, b*, and *n* contribute little to eqs. 20 and 21, which, as *N* increases, are dominated by products of 16*N* ^2^ and reduction factors due to consanguinity.

Recall that an exponentially distributed random variable with mean *λ* has variance *λ*^2^. Taking the ratio of eq. 21 and the square of eq. 15, as *N* → ∞, Var[*V*]*/* 𝔼[*V*]^2^ → 1. This relationship suggests *V* might be exponentially distributed in the *N* → ∞ limit. Considering eqs. 20 and 13, we find 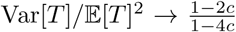, so for *c* > 0, *T* is not exponentially distributed in the limit.

## 4 Limiting distribution of coalescence times

With the exact variance established, we now examine the full distribution of coalescence times under the model, in the limit of large *N*. As a model in which two alleles can either coalesce rapidly due to consanguinity or reenter the ancestral process, the model has two time scales on which coalescence can take place. It is therefore suited to use of the separation-of-time-scales approach of MÖhle (1998b). We next review this approach as background to analysis of coalescence times in our consanguinity models.

### 4.1 Separation-of-time-scales approach

In the separation-of-time-scales approach, we can describe the ancestral process by a single-generation transition matrix Π_*N*_, for transitions between states permissible for a pair of alleles. MÖhle (1998b) derived the limiting distribution of coalescence times in cases where the transition matrix can be written as

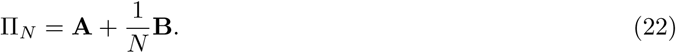

This approach splits the matrix Π_*N*_ into “fast” transitions in **A** that occur with rate 𝒪 (1) and “slow” transitions in 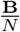 that occur with rate 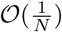. In other words, matrix **A** describes the rapid coalescent events that occur due to the part of the process that occurs on a relatively fast time scale, and matrix **B** includes the slower events that occur on a time scale proportional to *N*.

Möhle showed that as *N* → ∞, in comparison to the slow time scale of **B**, the fast process of **A** appears instantaneous and is characterized by the equilibrium

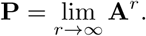

With time *t* scaled in units of *N* generations, Π_*N*_ converges weakly to a continuous-time process, such that

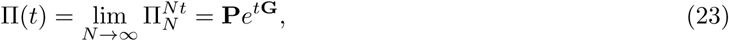

where the rate matrix is **G**=**PBP**.

In the following sections, we apply Möhle’s results to our models. We write the transition matrix Π_*N*_ for our population with consanguinity and decompose Π_*N*_ into **A** and **B**. Next we find the equilibrium **P**, and we use **P** and **B** to compute rate matrix **G**. Finally, we derive the exponential (eq. 23) to find the limiting distribution of coalescence times.

### 4.2 Sib mating

Recall that our sib mating model has *N* mating pairs, a fraction *c*_0_ of which are siblings. In this model, two alleles can be in four states: state 0, coalescence; state 1, within an individual; state 2, in two individuals in a mating pair; and state 3, in two individuals in separate mating pairs. If two alleles are in state 0, then they remain coalesced with probability 1. If two alleles are in an individual (state 1) then in the previous generation they are in two individuals in a mating pair with probability 1 (state 2). If the two alleles are in state 2, then with probability *c*_0_, the mating pair is a sib mating pair, and in the previous generation, the alleles transition to states 0, 1, and 2 with probabilities *c*_0_*/*4, *c*_0_*/*4, and *c*_0_*/*2, respectively. If the two alleles are not in a sib mating pair, then they transition to state 3 in the previous generation. Similarly, if two alleles are in state 3, then with probability 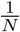 the individuals are siblings, and the alleles can transition to states 0, 1, and 2 with probabilities 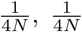, and 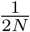, respectively. With probability 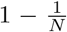, the two individuals are not siblings, and the alleles remain in state 3. These cases give the transition matrix

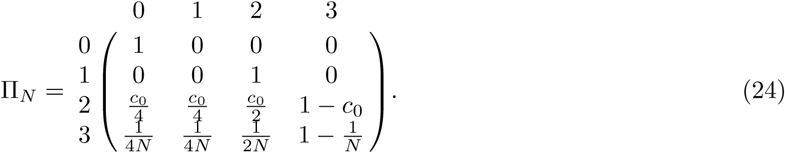

Decomposing Π_*N*_ (eq. 24) into fast and slow transitions as in eq. 22, we can write matrices **A** and **B** as

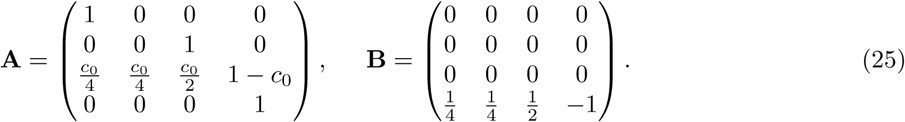

To find Π(*t*) we first derive the limit of matrix **A**. This computation, performed in Appendix A, gives

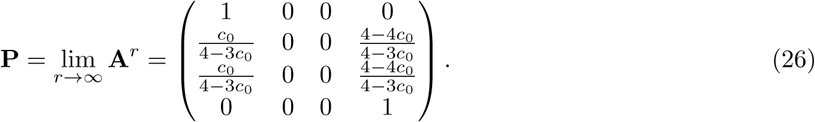

Next, we compute the rate matrix **G** by taking the product

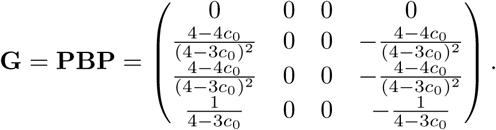

Finally, we apply eq. 23, computing the matrix exponential **P***e*^*t***G**^. Converting *t* back to units of generations, we evaluate 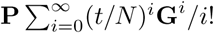,

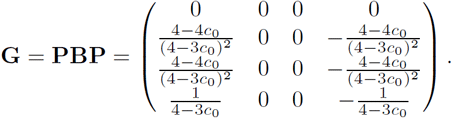

From the bottom three rows of the first column of Π(*t*), corresponding to states 1, 2, and 3, respectively, we extract the limiting cumulative distribution functions for *T, U*, and *V* :

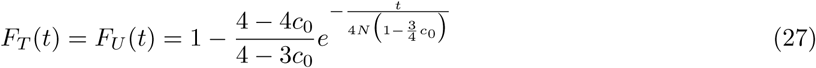

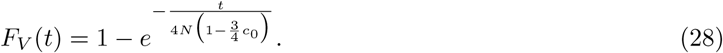

We immediately observe in eq. 28 that *V* is exponentially distributed with mean 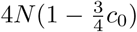, so that the coalescence time of two alleles in two individuals in separate mating pairs is distributed identically to that of two alleles in a haploid population of size 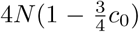. With no consanguinity, *c*_0_ → 0 as *N* → ∞, *F*_*T*_ (*t*) = *F*_*V*_ (*t*), and *T, U*, and *V* are all distributed identically to the coalescence time for two alleles in a haploid population of size 4*N*.

For *c*_0_ > 0, *T* and *U* are not exponentially distributed. We have 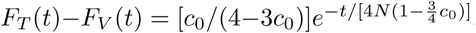, so *F*_*T*_ (*t*) > *F*_*V*_ (*t*) and the probability that two alleles in an individual coalesce by *t* generations ago exceeds the corresponding probability for two alleles in separate mating pairs. As *c*_0_ increases to 1 at fixed *N* and *t, F*_*T*_ (*t*) − *F*_*V*_ (*t*) increases; for fixed *N* and *c*_0_, the difference is largest at *t* = 0, decreasing to 0 as *t* increases.

From eqs. 27 and 28, noting that for a random variable *X* ≥ 0 with cumulative distribution function 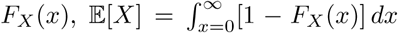 and 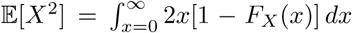, we can compute the mean and variance of the limiting distributions of *T* and *V* as

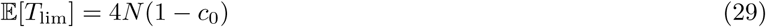

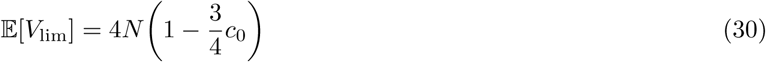

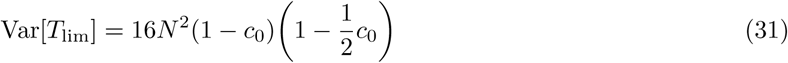

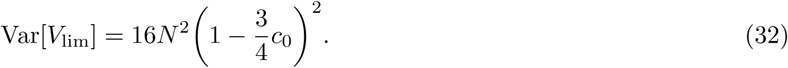

The differences between the large-*N* limiting values and the exact solutions in eqs. 1, 3, 11, and 12 are

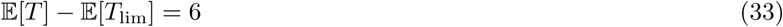

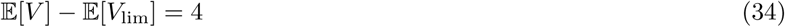

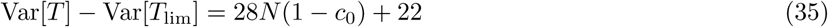

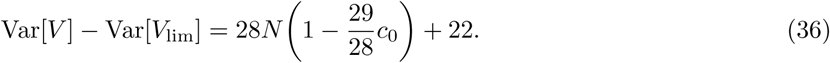

For large *N*, the differences in eqs. 33-36 are negligible in comparison to the exact means and variances.

Eqs. 27 and 28 are plotted in Figure 3. In Figure 3A, we observe that as *c*_0_ increases, the probability of instantaneous coalescence increases. This probability is maximized for *c*_0_ = 1, where *F*_*T*_ (*t*) = 1, implying that all pairs of alleles coalesce quickly due to consanguinity. For *V*, we observe in Figure 3B that increased consanguinity decreases mean coalescence time 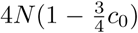, and the distribution behaves like a random-mating population with a reduced size. Compared to a haploid population of size 4*N*, for both *T* and *V*, as consanguinity increases, probability density is shifted towards zero, away from ancient coalescence times to more recent coalescence times.

**Figure 3:**
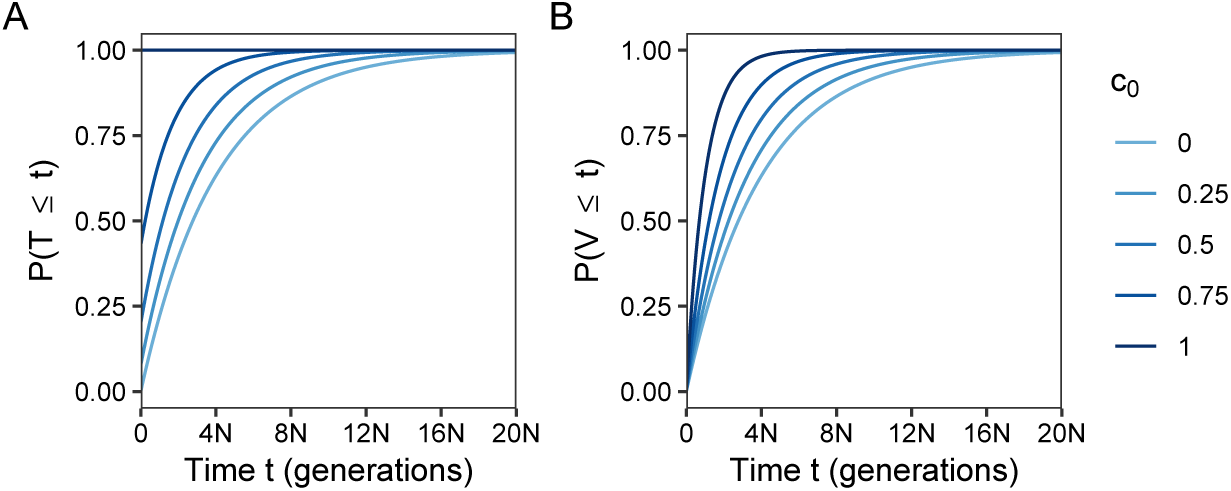
The cumulative distributions of coalescence times within (*T*) and between (*V*) individuals as functions of generations *t* and the fraction *c*_0_ of sib mating pairs. **(A)** ℙ (*T* ≤ *t*), eq. 27. **(B)** ℙ (*V* ≤ *t*), eq. 28.

### 4.3 Superposition of multiple mating levels

We now generalize the separation-of-time-scales approach to allow a superposition of mating levels. Recall that under the superposition, *i*th-cousin mating is permitted for each *i* from 0 to *n*, where *n* is the degree of the most distant permissible cousin relationship. For each *i*, let *c*_*i*_ be the fraction of *i*th-cousin mating pairs each generation, with 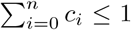. As before, we assume that *i*th-cousin mating pairs are distinct sets for distinct *i*, such that two individuals in a mating pair cannot, for example, be both first and third cousins.

We derive the single-generation transition matrix Π_*N*_. We have states 0, 1, and 3 as before, and transitions from these states are the same as under sib mating. For state 2, if two alleles are in two individuals in a mating pair in the current generation, then for each *i*, 0 ≤ *i* ≤ *n*, with probability *c*_*i*_*/*4^*i*^, the two individuals are *i*th cousins and the alleles were inherited from the shared ancestral mating pair *i* + 1 generations in the past. For each *i*, 0 ≤ *i* ≤ *n*, there exists a state, which we call 2_*i*_, that is visited *i* generations back in time from the current generation. For example, the state that we termed state 2 under sib mating we now call 2_0_, because with probability *c*_0_, the two individuals in the mating pair are 0th cousins (sibs) who share an ancestral mating pair 1 generation in the past. If two alleles have reached state 2_*i*_, then they have not yet coalesced, and the two individuals in the current generation are not more closely related than *i*th cousins. If *c*_*i*_ > 0, then the individuals might be *i*th cousins and two alleles in state 2_*i*_ can be inherited from the shared ancestral mating pair in the next generation back in time from state 2_*i*_ (Figure 4).

**Figure 4:**
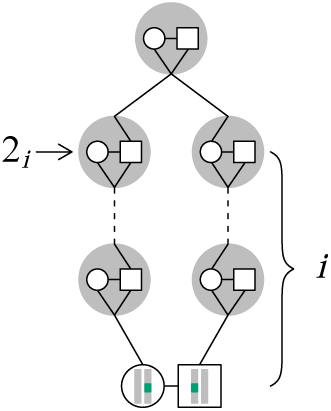
New states in the superposition model. Two alleles in two individuals who are in an *i*th cousin mating pair and have no closer relationship, *i* generations ago, are in separate mating pairs. We term their state *i* generations ago 2_*i*_. In the next generation back, the alleles might be in the shared ancestral mating pair, as shown. Here, *i* = 2.

For convenience, for each *i*, 0 ≤ *i* ≤ *n*, we define *k*_*i*_ as the probability that two alleles in two individuals in a mating pair coalesce due to consanguinity in at most *i* + 1 generations, so that the individuals are *i*th cousins or more closely related,

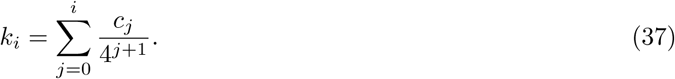

We denote *k*_−1_ = 0. The probability that the alleles in two individuals in a mating pair *reach* a shared ancestral pair in at most *i* + 1 generations is 4*k*_*i*_; the conditional probability that they coalesce in that pair given that they have reached it is 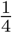. Then 1 − 4*k*_*i*_ is the probability that two alleles in two individuals in a mating pair are in a pair with relationship more distant than *i*th cousins; it is the probability that the individuals have no shared ancestor up to and including *i* + 1 generations back from the present.

We next define *x*_*i*_ for 0 ≤ *i* ≤ *n* as the conditional probability that two alleles in a mating pair are in an *i*th-cousin mating pair and that they coalesce in the shared ancestral pair *i* + 1 generations back, given that they have no shared ancestor *i* generations back or more recent. The probability that two alleles in a mating pair are in an *i*th-cousin pair is *c*_*i*_, the probability that they coalesce in the shared ancestral pair is 1*/*4^*i*+1^, and the probability that they have no shared ancestor *i* generations back or more recent is 1 − 4*k*_*i*−1_. Hence,

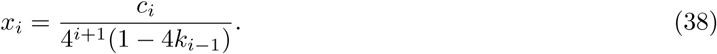

If two alleles are in state 2_*i*_, then they are in lineages ancestral to two individuals who are not more closely related than *i*th cousins, and they have not coalesced. Two alleles in state 2_*i*_ have four possible transitions: with probability *x*_*i*_, they coalesce (state 0); with probability *x*_*i*_, they are inherited from the same individual in the ancestral mating pair but do not coalesce (state 1); with probability 2*x*_*i*_, they are inherited from two individuals in the ancestral mating pair (state 2_0_). With probability 1 − 4*x*_*i*_, the two alleles were not inherited from a shared ancestor *i* + 1 generations ago, so the individuals in the current generation are not more closely related than (*i* + 1)th cousins, and the alleles transition to state 2_*i*+1_. If the alleles are in state 2_*n*_, then they transition to states 0, 1, and 2_0_, as seen for states 2_*i*_, *i < n*, but the fourth transition is to two individuals in separate mating pairs (state 3), with probability 1 − 4*x*_*n*_.

Combining these cases gives the transition matrix Π_*N*_ over states 0, 1, 3, and 2_*i*_ for 0 ≤ *i* ≤ *n*:

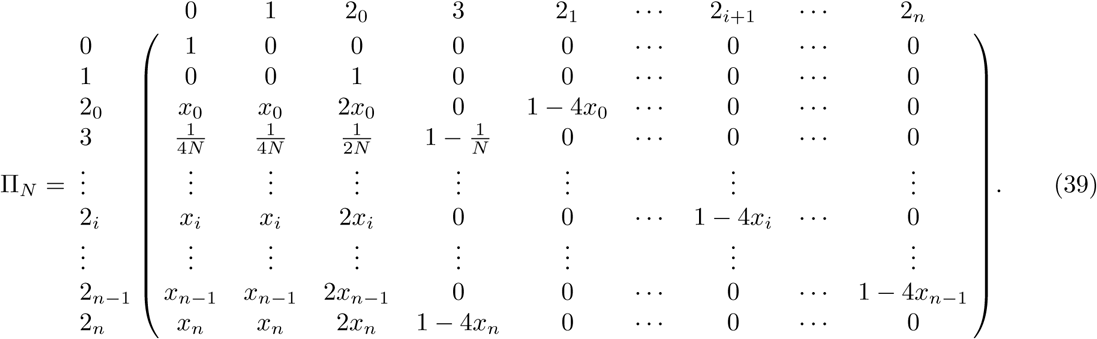

Π_*N*_ decomposes into the 𝒪 (1) transitions in matrix **A**,

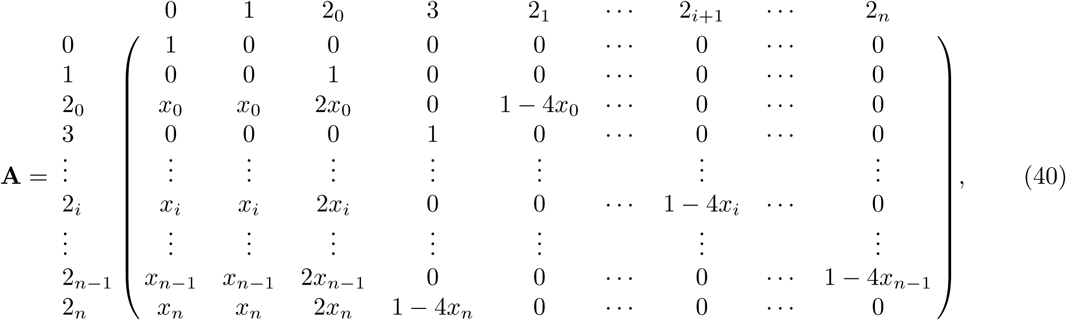

and the 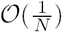 transitions in matrix **B**,

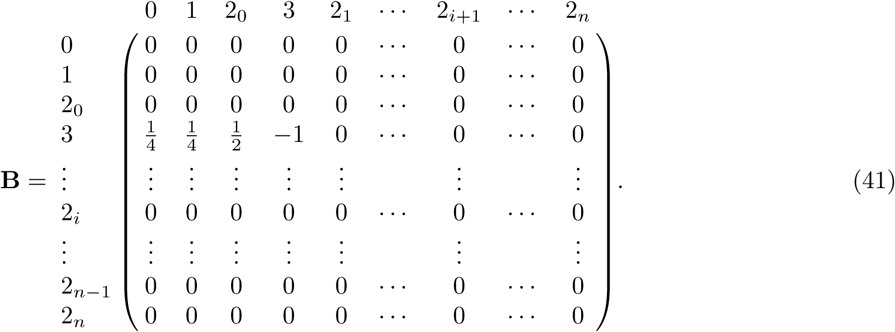

We derive the equilibrium of **A** in Appendix B. We obtain

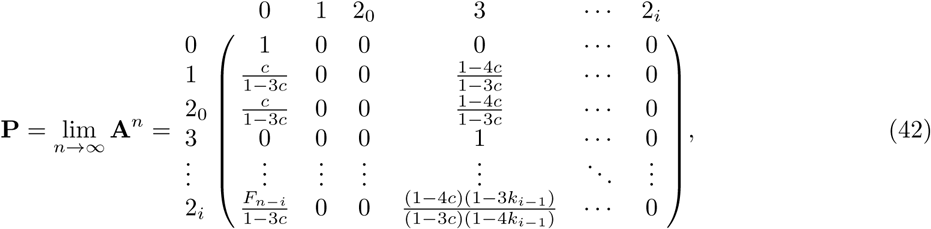

where *F*_*n*−*i*_ is described in Appendix B (**P** has the same dimension as **A** and **B**, but for convenience we do not expand down to rows 2_*n*−1_ and 2_*n*_ in **P**, as they follow the form of row 2_*i*_). Because **A** is an absorbing matrix with absorbing states 0 and 3, the only nonzero entries in **P** are in columns 0 and 3. Note that at equilibrium, 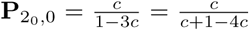. The numerator of this fraction is the probability that two alleles in a mating pair coalesce rapidly due to consanguinity, *c*, and the denominator is the sum of this quantity and the probability the two alleles are not inherited through the consanguineous pedigree, 1 − 4*c*. Note that for the sib mating case, *c* = *c*_0_*/*4, and *c/*(1 − 3*c*) becomes *c*_0_*/*(4 − 3*c*_0_), as seen in eq. 26.

Next we take the product **PBP** (Appendix B) to find matrix **G**:

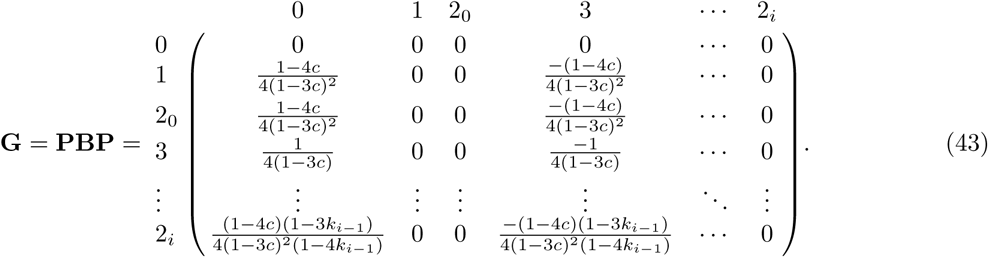

Lastly, to derive Π(*t*), we compute the exponential *e*^*t***G**^ (Appendix B) and take the product **P***e*^*t***G**^. With *t* measured in units of generations, we obtain

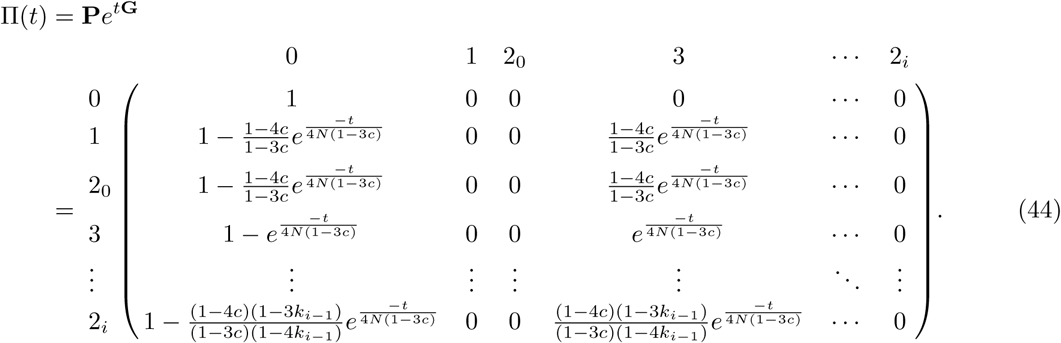

As we observed for **P**, the only nonzero entries of Π(*t*) are in columns 0 and 3.

From column 0 of Π(*t*), examining the rows for states 1, 2_0_, and 3, respectively, we have the limiting cumulative distributions for coalescence times *T, U*, and *V* :

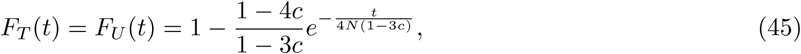

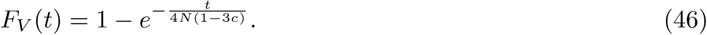

We immediately observe that for the sib mating case of *c* = *c*_0_*/*4, eqs. 45 and 46 reduce to eqs. 27 and 28, respectively. In the limit, *T* and *U* are identically distributed but not exponential. The limiting *V* is exponentially distributed with mean 4*N* (1 − 3*c*); the coalescence time of two alleles in two individuals in separate mating pairs is therefore identically distributed with that of two alleles in a haploid population of size 4*N* (1 − 3*c*). Consanguinity reduces effective population size compared to random mating, with the reduction dependent on the kinship coefficient *c* of a randomly chosen mating pair.

The means and variances of the limiting distributions are

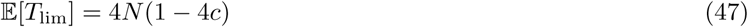

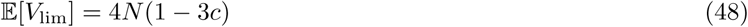

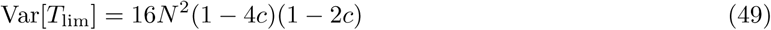

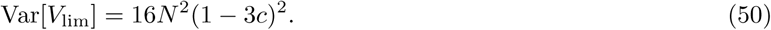

Considering eqs. 13 and 15, the differences between the exact and limiting means of *T* and *V* are

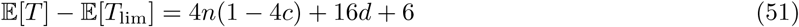

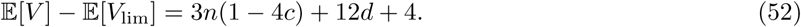

The exact means exceed the limiting means for *c* > 0. Recall from Section 3.2 that 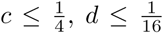, and 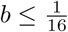. Then for *n* ≪ *N*, eqs. 51 and 52 contribute little to 𝔼[*T*] and 𝔼[*V*].

For the differences between the variances, we have

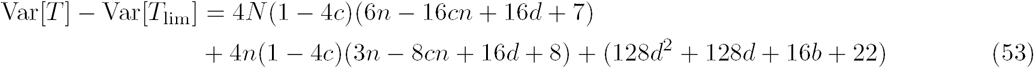

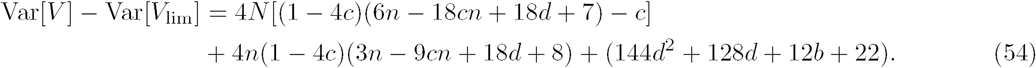

For large *N*, because Var[*T*] and Var[*V*] are 𝒪 (*N* ^2^), the differences contribute relatively little in relation to the magnitudes of the variances. Finally recall that if *c* = *c*_0_*/*4, then *d* = 0, *b* = 0, and *n* = 0, and the quantities in eqs. 47-54 reduce to those for sib mating, eqs. 29-36.

For *c* = 0, there is no consanguinity, *F*_*T*_ (*t*) = *F*_*V*_ (*t*), and *T* and *V* are both exponentially distributed. For *c* > 0, the difference 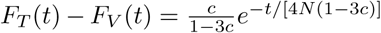 is positive, and *F*_*T*_ (*t*) > *F*_*V*_ (*t*). The probability that two alleles within an individual have coalesced by time *t* is greater than or equal to the probability for two alleles in separate individuals. As *c* increases to 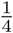, the difference increases; for fixed *c*, as *t* increases, the difference approaches zero, so that it is greatest for recent coalescence times.

## 5 Simulated distributions from the Markov chain

### 5.1 Simulation method

To examine the extent to which the limiting distributions of *T* and *V* accord with the exact distributions, we simulate pairwise coalescence times from the exact Markov chain (eq. 39). For both *T* and *V*, we consider a range of values of the number of mating pairs, *N* = 10, 20, 50, 100, 200, 500, 1000; the degree of cousin relationship, *n* = 0, 1, 2, 3, 4, 5; and the consanguinity rate, *c*_*n*_ = 0, 0.1, 0.2, 0.5, 0.75. For simplicity, we consider only one type of cousin relationship at a time. For each of the two random variables, *T* and *V*, and each set of parameter values {*N, n, c*_*n*_ }, we simulated 10^6^ pairwise coalescence times.

To compare limiting distributions of coalescence times (eqs. 45 and 46) and simulated exact distributions, we compute a chi-square test statistic. We divide the limiting cumulative distribution functions into intervals and count occurrences of simulated coalescence times within those intervals. For *V*, we divide the limiting function into 50 intervals of equal probability 0.02. The limiting function for *T* is nonzero at *t* = 0; if the probability at 0 is 0.02 or greater, then the first interval is assigned size *f*_*T*_ (0), and the remaining probability is divided into *q* = *l*(1 − *f*_*T*_ (0))*/*0.02*J* intervals, each with size (1 − *f*_*T*_ (0))*/q* ≥ 0.02.

### 5.2 Simulation results

The chi-square test statistics appear in Figure 5. Within each panel, we see that as *N* increases, the statistic generally decreases and then levels off, suggesting that increased population size improves the agreement between the exact and limiting distributions. This result accords with the fact that the limiting distribution is a large-*N* approximation, expected to more closely approximate the exact distribution as *N* increases.

**Figure 5:**
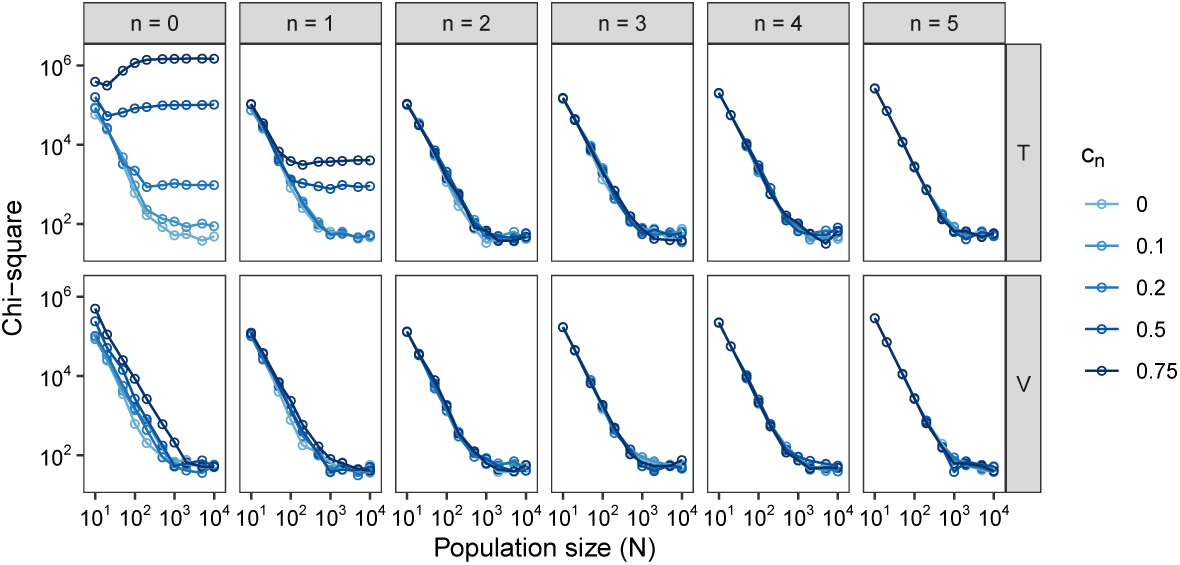
Values of the chi-square test statistic comparing the limiting distributions for *T* and *V*, eqs. 45 and 46, and the simulated exact distributions. The plots consider a range of values for the number of mating pairs (*N*), the consanguinity rate (*c*_*n*_), and the degree of cousin relationship (*n*).

Considering the panels from left to right, as *n* increases past *n* = 1, the agreement is similar at different levels of consanguinity *c*_*n*_. Thus, for relationships at the level of second or more distant cousins, the number of mating pairs *N* is the most important determinant of the agreement of the limiting and exact distributions. Examining the bottom row of Figure 5, for the random variable *V*, although for fixed *N*, the agreement is somewhat reduced at greater *c*_*n*_, a key role for *N* is also observed for *n* = 0 and *n* = 1.

In the top row of Figure 5, for random variable *T*, we see that for *n* = 0 and *n* = 1, at high *c*_*n*_, agreement between the limiting and exact distributions is relatively poor. In these cases, the probability of immediate coalescence in time 0 is larger in the limiting distribution. In the limiting distribution, with *c* = *c*_*n*_*/*4^*n*+1^, this probability is *c/*(1 − 3*c*) for *f*_*T*_ (0), and in the exact distribution, coalescence due to consanguinity has probability *c*. For large *c*, as occurs for large *c*_0_ or *c*_1_, *c/*(1 − 3*c*) ≉*c*.

In Figure 6, we more closely examine the effect of population size on the agreement between the limiting and simulated exact distributions. Over a range of population sizes, with *n* = 1 and a first-cousin relationship *c*_1_ = 0.2, we plot the cumulative distribution functions of *T* and *V* for the first 4*N* generations. Considering plots from left to right, for both *T* and *V*, as *N* increases, the limiting distribution more closely matches the simulated exact distribution. In the small-*N* plots with *N* = 10, we can observe that the limiting distribution begins at *t* = 0 with a higher cumulative probability, and that this excess persists as *t* increases.

**Figure 6:**
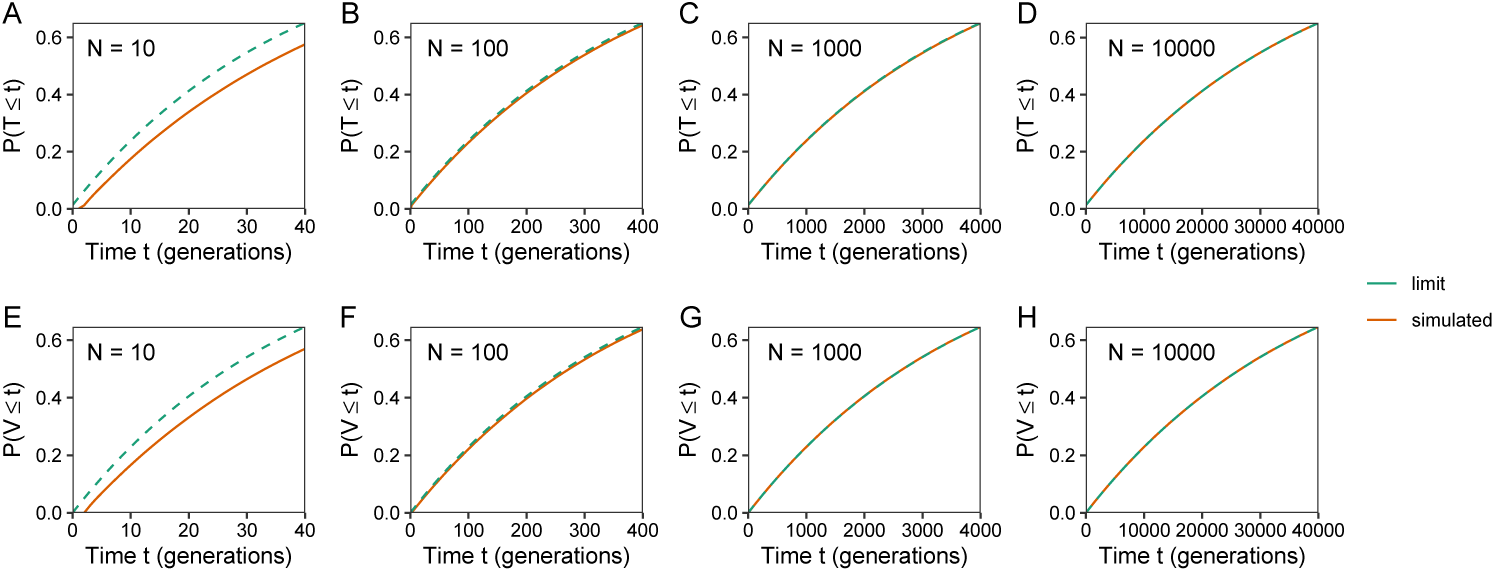
Limiting cumulative distribution functions for *T* and *V*, eqs. 45 and 46, and simulated exact cumulative distributions, for 4*N* generations. The plots consider a range of values for the number of mating pairs (*N*), fixing the degree of cousin relationship at *n* = 1 and the consanguinity rate at *c*_1_ = 0.2. (A) *T, N* = 10. (B) *T, N* = 100. (C) *T, N* = 1000. (D) *T, N* = 10000. (E) *V, N* = 10. (F) *V, N* = 100. (G) *V, N* = 1000. (H) *V, N* = 10000.

In Wakeley *et al.* (2012), the disagreement between coalescence time distributions for two models, a pedigree model with *N* individuals and the Kingman coalescent, was greatest in the most recent log_2_(*N*) generations. As our consanguinity models are similar to the pedigree model in that consanguinity influences the probability of rapid coalescence, we next examined the agreement of the limiting and simulated exact coalescent time distributions in the most recent generations. With the same parameter values as in Figure 6, Figure 7 focuses on the first 25 generations. For *T*, in Figure 7A-D, a difference occurs between the limiting and simulated exact distributions during the most recent generations, as the limiting distribution has a point mass at *t* = 0. For *V*, in Figure 7E-H, the limiting distribution does not have a point mass at *t* = 0, and the distributions differ by an amount that is approximately constant over the first 25 generations.

**Figure 7:**
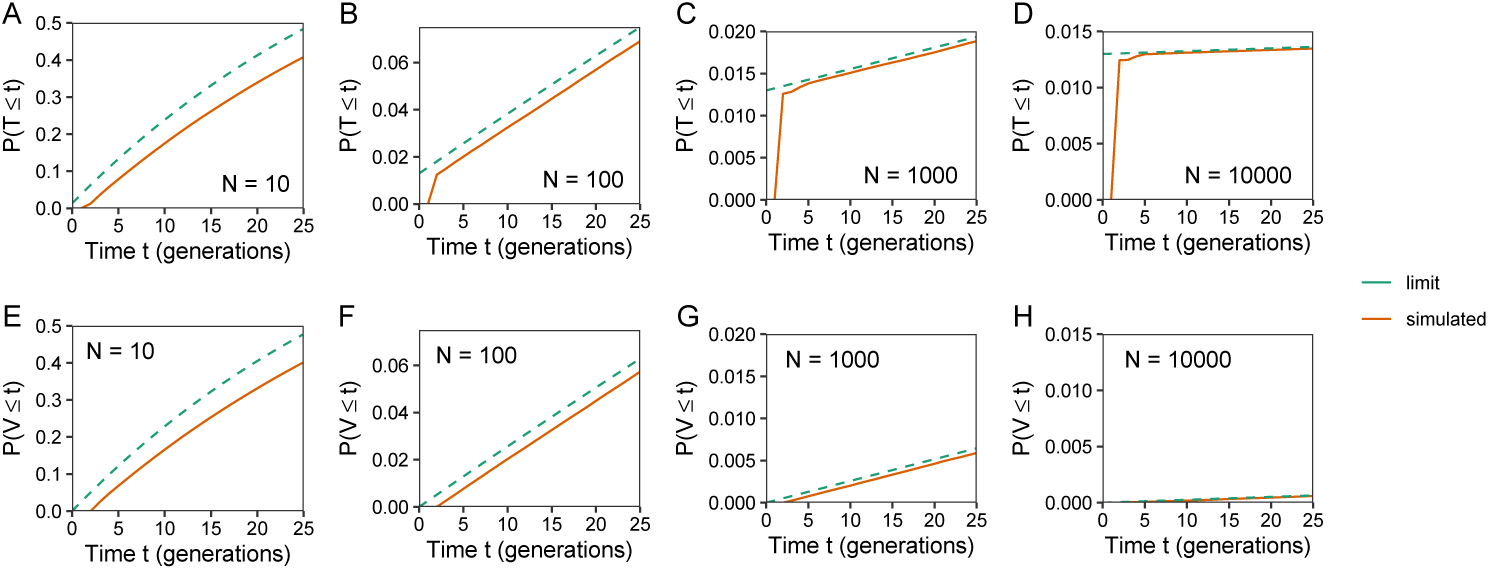
Limiting cumulative distribution functions for *T* and *V*, eqs. 45 and 46, and simulated exact cumulative distributions, for 25 generations. The plots consider a range of values for the number of mating pairs (*N*), fixing the degree of cousin relationship at *n* = 1 and the consanguinity rate at *c*_1_ = 0.2. (A) *T, N* = 10. (B) *T, N* = 100. (C) *T, N* = 1000. (D) *T, N* = 10000. (E) *V, N* = 10. (F) *V, N* = 100. (G) *V, N* = 1000. (H) *V, N* = 10000.

## 6 Discussion

Building on a study of mean coalescence times under consanguinity in a diploid model with *N* mating pairs, we have expanded the analysis to examine full coalescence time distributions. Under sib mating, we calculated the exact variance of coalescence times for two alleles within an individual and two alleles in separate individuals (eqs. 11 and 12), and we generalized the result to a superposition of multiple levels of cousin mating (eqs. 20 and 21). Using separation of time scales to examine “fast” coalescence by consanguinity and “slow” coalescence in the general population, we derived the large-*N* limiting distribution of pairwise coalescence times for two alleles within an individual and two alleles in separate individuals, in both the sib mating (eqs. 27 and 28) and superposition models (eqs. 45 and 46). As *N* increases, distributions simulated from the exact Markov chain approach the limiting distributions (Figures 5-7).

Previously (Severson *et al.*, 2019), we showed that increased consanguinity reduces mean pairwise coalescence times both within and between individuals, with a stronger effect for two alleles within an individual. In each of several models, we found that the reduction factor could be written in terms of the kinship coefficient *c* of a randomly chosen mating pair. Here, by deriving limiting distributions of coalescence times, we can further explain the earlier result. In particular, for two alleles in separate individuals, limiting coalescence times are distributed with pairwise coalescence times as in a haploid population of size 4*N* (1 − 3*c*). For two alleles within an individual, the distribution is a mixture of this effective size reduction and instantaneous coalescence with probability 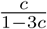. Increasing consanguinity reduces the coalescent effective size (Sjodin *et al.*, 2005) of the “slow” process; in the large-*N* limit, if rapid coalescence due to consanguinity does not occur, then coalescence follows the standard haploid model with the reduced population size.

The view of our model as having rapid coalescence due to consanguinity followed by coalescence mimicking a standard haploid population aligns with similar results for other phenomena that permit the separation-of-time-scales approach (Wakeley, 2009, chapter 6). Related models consider partial selfing (Nordborg and Donnelly, 1997; MÖhle, 1998b; Nordborg and Krone, 2002), two sexes (MÖhle, 1998a; Nordborg and Krone, 2002), stage structure (Nordborg and Krone, 2002), many-demes migration (Wakeley, 2001, 2004; Eldon and Wakeley, 2009), and combinations of factors, as in an analysis of two sexes, sex chromosomes, and migration (Ramachandran *et al.*, 2008).

The parallel is most natural for partial selfing. Following Nordborg and Krone (2002), consider a diploid population of 2*N* individuals, in which the probability that two alleles within an individual coalesce in the previous generation is 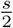, where *s* is the selfing rate—the fraction of individuals for whom the same parent provides both of their genomic copies. In the selfing model, 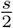 is the probability of immediate coalescence in one generation, and 1 − *s* is the probability that a pair of alleles “escapes” from the rapid time scale of coalescence by selfing. In the large-*N* limit, the probability of rapid coalescence is 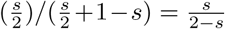. This result has a similar structure to our large-*N* result that the probability of rapid coalescence by consanguinity for two alleles in a mating pair is 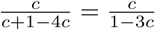, where *c* is the probability of coalescence by consanguinity during the first *n* generations and 1 − 4*c* is the probability of escape into the slow process. In both cases, a probability exists that the alleles return to the initial configuration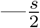 for the selfing model and 3*c* for the consanguinity model—with the chance to either coalesce in the fast process or escape to the slow one.

Our interest in studying coalescence time in a consanguinity model has been motivated by the link between coalescence times and lengths of genomic segments shared identically by descent, with the random variable *T* connecting to ROH within individuals, and *V* connecting to IBD tracts between genomes in separate individuals chosen at random in a population (Severson *et al.*, 2019). For a pair of genomes, the random length of the segment shared identically by descent around a locus is inversely related to the random pairwise coalescence time at the focal locus, with recombination acting to shorten the shared fragment. In our previous work (Severson *et al.*, 2019), we used the inverse relationship between the mean coalescence time and shared fragment length to provide qualitative results on trends in ROH and IBD sharing in relation to consanguinity. Under a recombination model, the distribution of the shared fragment length can be obtained from the full distribution of pairwise coalescence times (Palamara *et al.*, 2012; Carmi *et al.*, 2013, 2014). As we have now obtained the limiting distributions of pairwise coalescence times, both within and between individuals, it will now be possible to deepen empirical analyses of the effect of consanguinity on patterns in shared genomic segments.

## Acknowledgments

We acknowledge support from National Institutes of Health grant R01 HG005855, United States–Israel Binational Science Foundation grant 2017024, and a National Science Foundation Graduate Research Fellowship.

## Appendix A. Sib mating

For sib mating, this appendix provides details of the computation of matrix **P**, the *r* → ∞ limit of **A**^*r*^. The matrix **A** appears in eq. 25. To find the desired limit, note that **A** is an absorbing matrix with absorbing states 0 and 3. Recall that for an absorbing matrix **D** with form

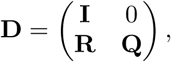

and with fundamental matrix **N** = (**I** − **Q**)^−1^, the limit of **D**^*r*^ is given by

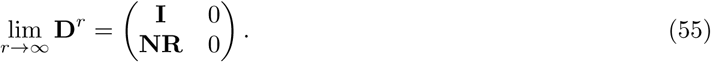

We rearrange **A** to match the form of **D** by permuting rows and columns to obtain permuted matrix **A**^∗^:

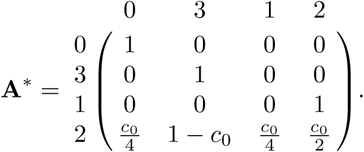

Now we can read **R** and **Q** as

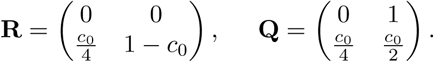

Next we compute the fundamental matrix **N**:

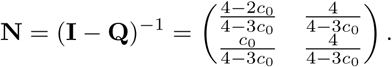

We find the product **NR**:

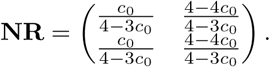

Following eq. 55, we have the desired limit,

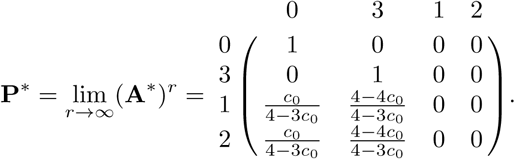

Permuting the columns and rows again, we obtain eq. 26.

## Appendix B. Superposition of multiple mating levels

This appendix provides details of the separation-of-time-scales computations in the case of a superposition of mating levels. We begin with some lemmas.

### Two lemmas

We recall *k*_*i*_ and *x*_*i*_ from eqs. 37 and 38. First, we will need a recursion *F*, defined by *F*_0_ = *x*_*n*_ and *F*_*m*_ = *x*_*n*−*m*_ + (1 − 4*x*_*n*−*m*_)*F*_*m*−1_ for *m* ≥ 1.

#### Lemma 1.

For *m* ≥ 1,

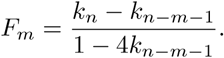

**Proof:** First consider the base case *m* = 0,

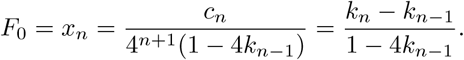

Next, we assume for induction that *F*_*m*−1_ = (*k*_*n*_ − *k*_*n*−*m*_)*/*(1 − 4*k*_*n*−*m*_). Then

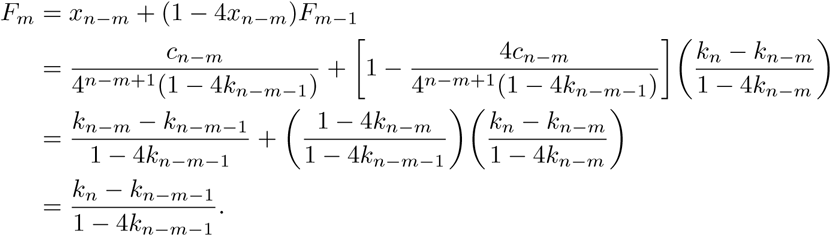

This completes the proof. □

#### Lemma 2.

For ℓ ≥ *j* ≥ 0,

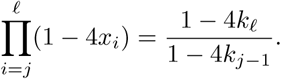

**Proof:** We use *c*_*i*_*/*4^*i*+1^ = *k*_*i*_ − *k*_*i*−1_ from eq. 37:

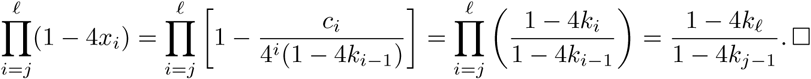

### The limiting matrix P

We follow the same method as in Appendix A. Again because **A** has two absorbing states, 0 and 3, we can derive the equilibrium matrix **P** with eq. 55. Permuting the columns and rows of **A** in eq. 40 from (0, 1, 2_0_, 3, *…*, 2_*i*_, *…*, 2_*n*_) to (0, 3, 1, 2_0_, *…*, 2_*i*_, *…*, 2_*n*_), **A**^∗^ has form

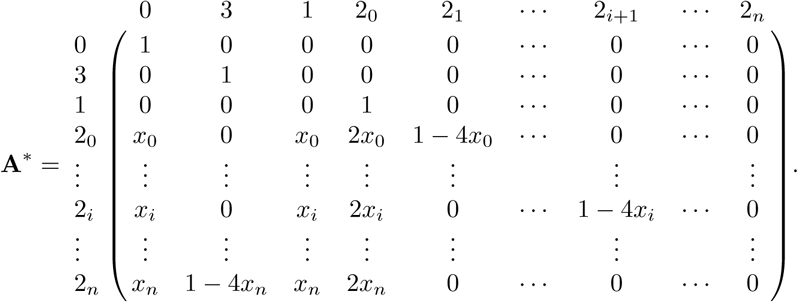

From **A**^∗^, we find **R** and **Q** as

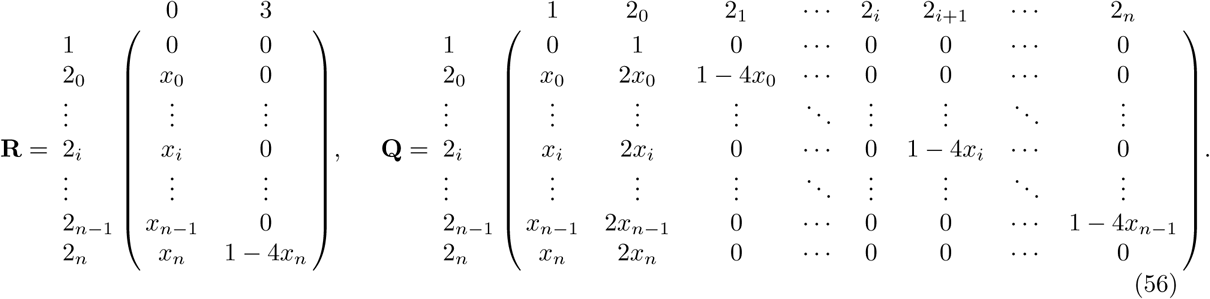

First, we find the fundamental matrix **N** = (**I** − **Q**)^−1^. We proceed by Gaussian elimination, beginning from the augmented matrix [(**I** − **Q**)| **I**] and proceeding to obtain (**I**|**N**). We write **M** = **I** − **Q**:

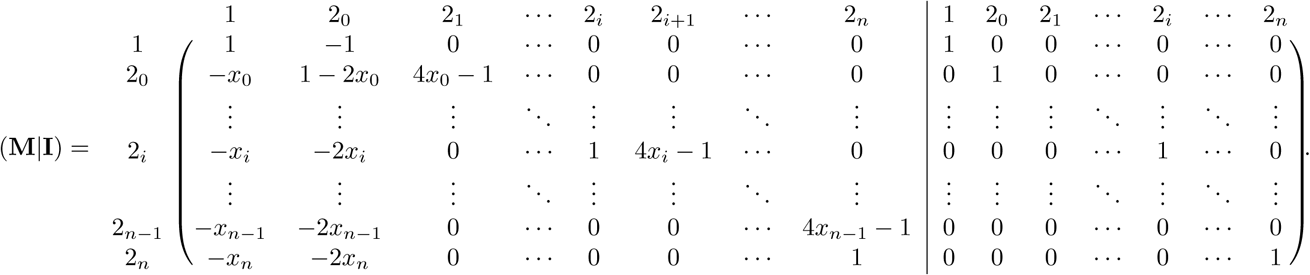

For convenience, we refer to rows and columns of **M** and **N** by their associated states, and continue to refer to the left and right components of the augmented matrix by **M** and **N**.

To begin the elimination, for each row 2_*i*_ we eliminate −*x*_*i*_ in the first column by adding *x*_*i*_ times row 1. As this step leaves a value of 4*x*_*n*−1_ − 1 in column 2_*n*_ of matrix **M**, we next add 1 − 4*x*_*n*−1_ times row 2_*n*_ to row 2_*n*−1_, obtaining

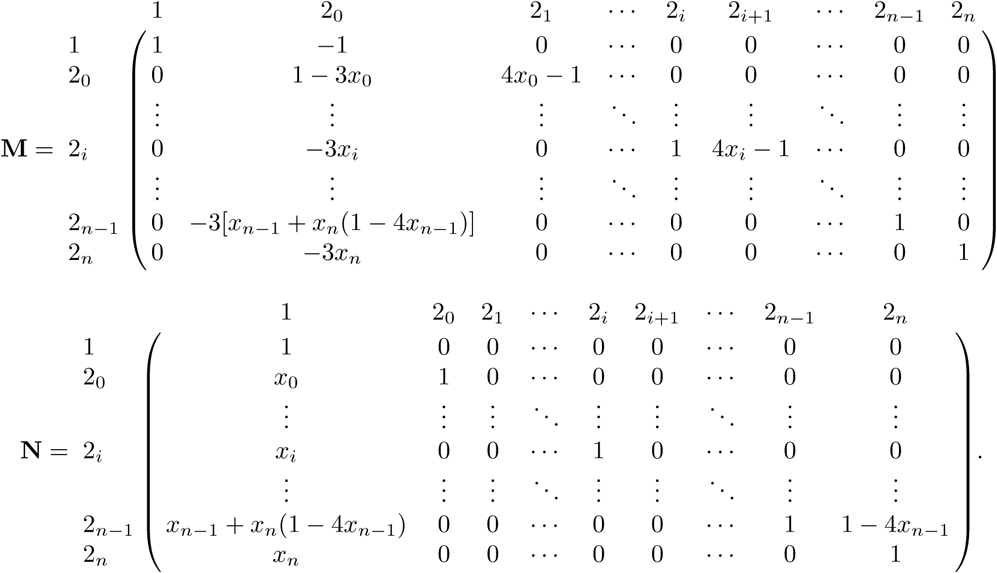

Notice that in **M**, for each row 2_*i*_ for *i* from 0 to *n*−2, the entry in column 2_*i*+1_ satisfies 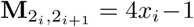. Now that we have eliminated 4*x*_*n*−1_ − 1 in row 2_*n*−1_, we repeat the same operation and use row 2_*n*−1_ to eliminate 4*x*_*n*−2_ − 1 in the row above, 2_*n*−2_. Specifically, we add (1 − 4*x*_*n*−2_) 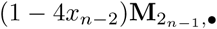 to 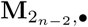 (where • indicates that here we consider row vectors). Decrementing *i* from *n* − 2 to 0, for each *i*, we perform this operation of adding to row 2_*i*_ the quantity 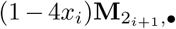 Repeatedly performing this operation produces a recursion in column 2_0_, and we have

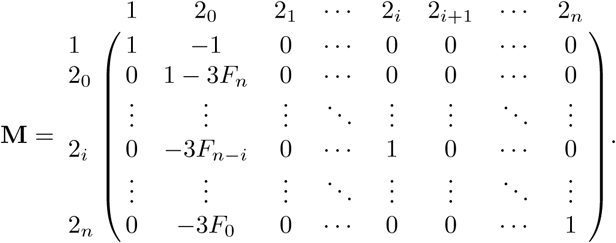

The operation of successively adding 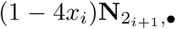 to 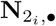 also produces the recursion *F* in column 1 of **N**. This operation creates increasing products of terms 1 − 4*x*_*i*_ in the upper right triangle of **N**. For an entry 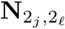, with ℓ > *j*, the entry is given by the product 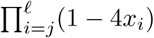. After completing this operation for all rows 2_*i*_ in **N**, 0 ≤ *i* ≤ *n* − 1, the matrix is

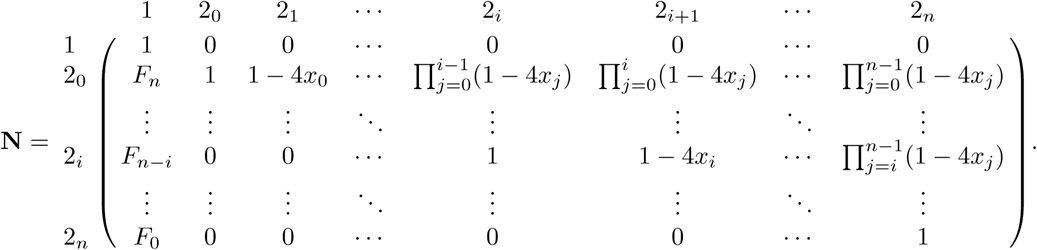

We can now simplify matrices **M** and **N** using Lemmas 1 and 2. First, by Lemma 1, *F*_*n*_ = (*k*_*n*_ − *k*_−1_)*/*(1 − 4*k*_−1_) = *k*_*n*_ = *c*, where *c*, defined in eq. 16, is the kinship coefficient of two individuals in a randomly chosen mating pair. Then we can rewrite **M** as

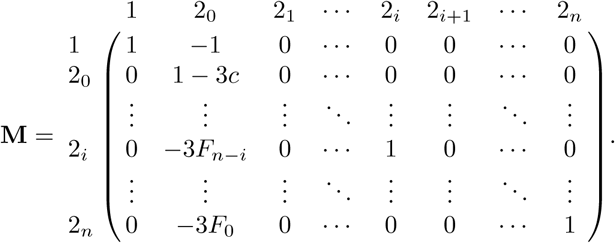

Using Lemma 2, we can simplify **N**:

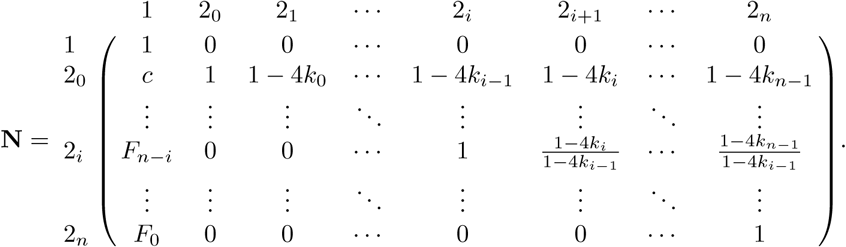

For the last elimination step in column 2_0_ of **M**, we first divide row 2_0_ of **M** and **N** by 1 − 3*c* and then add the resulting row 2_0_ to row 1. We obtain:

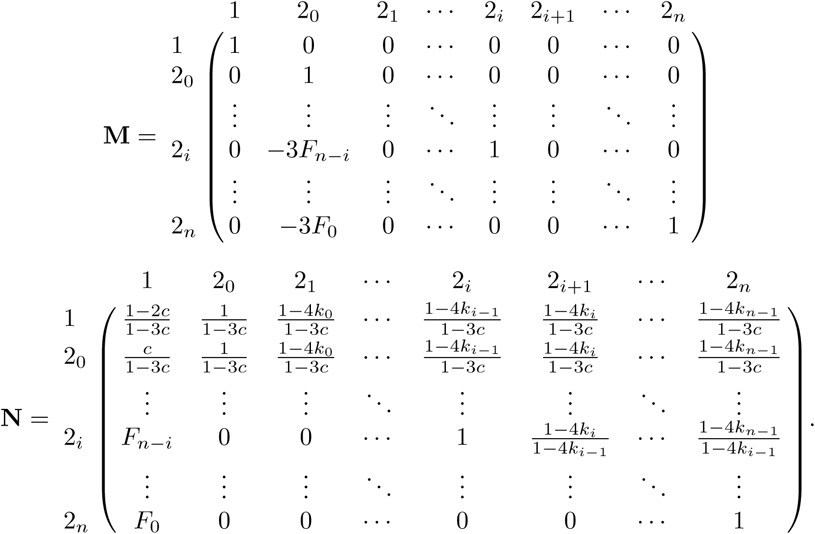

Next, for each remaining row 2_*i*_, 1 ≤ *i* ≤ *n*, in **M**, we add 3*F*_*n*−*i*_ times row 2_0_, which for **M** gives

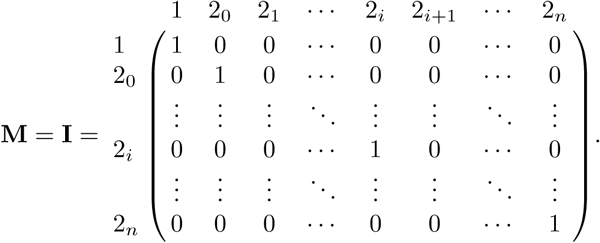

For **N**, this step produces the fundamental matrix **N** = (**I** − **Q**)^−1^, with form

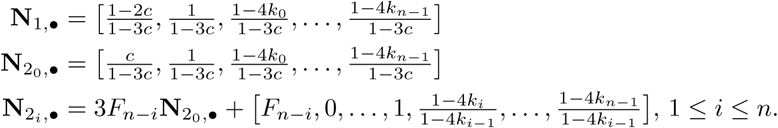

Now that we have derived the fundamental matrix **N**, we next find the product **NR**. From eq. 56,

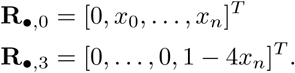

Because **R**_1,•_ = [0, 0] and rows **N**_1,•_ and 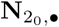 only differ at their first entry, we have dot products **N**_1,•_. 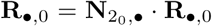 and 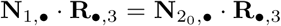. Hence, to complete the derivation of **NR**, it suffices to compute the dot products of **N**_1,•_ and 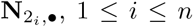, 1 ≤ *i* ≤ *n*, with **R**_•,0_ and **R**_•,3_. Using eqs. 37 and 38 and Lemma 1, these products are

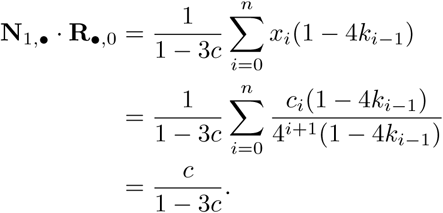

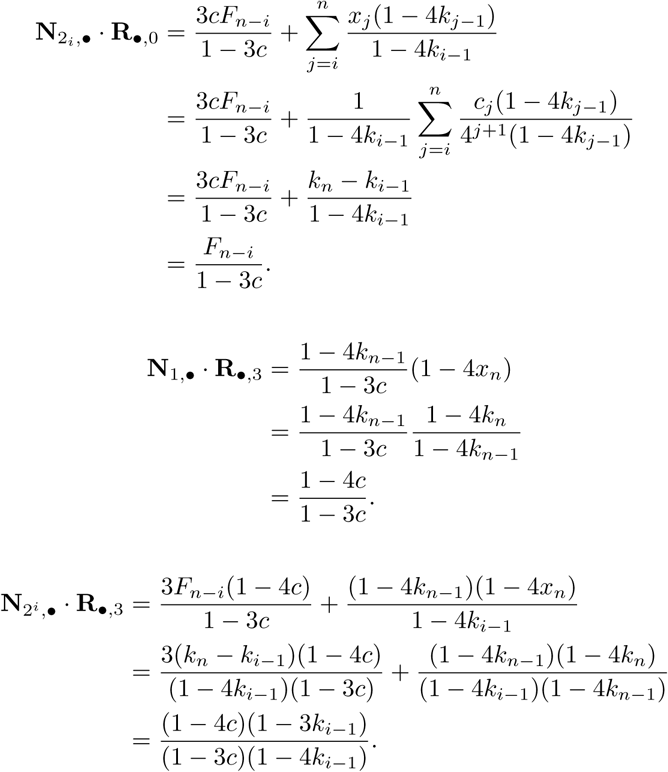

Combining these cases, we find

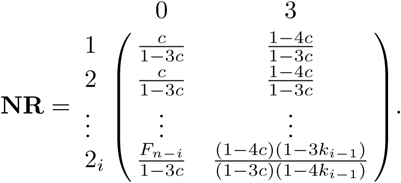

We have

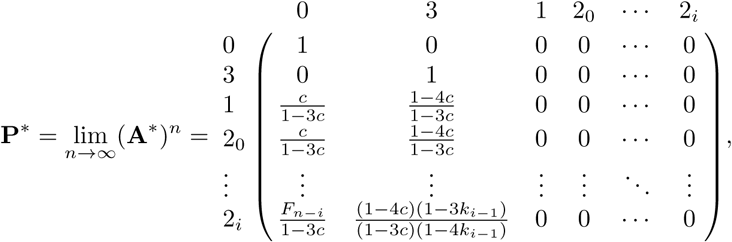

from which we obtain **P** in eq. 42 by permuting rows and columns.

### The generator matrix G

Here we derive the generator matrix **G=PBP**. Recall **B** from eq. 41. We first compute **BP**.

Because **B**_3,•_ is the only nonzero row of **B**, the only nonzero row of **BP** is (**BP**)_3,•_. Similarly, because columns **P**_•,0_ and **P**_•,3_ are the only nonzero columns of **P**, the only nonzero columns of **BP** are (**BP**)_•,0_ and (**BP**)_•,3_. Therefore the only nonzero entries of **BP** are (**BP**)_3,0_ and (**BP**)_3,3_:

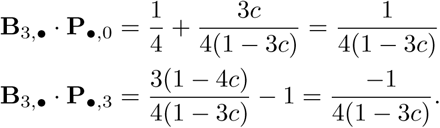

Hence, we have

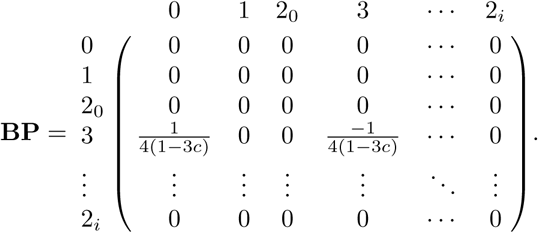

Next, for the product **PBP**, we note again that because only columns (**BP**)_•,0_ and (**BP**)_•,3_ are nonzero, the only nonzero columns of **PBP** are 0 and 3. Because the only nonzero elements of columns (**BP**)_•,0_ and (**BP**)_•,3_ are in row (**BP**)_3,•_, the entries in columns (**PBP**)_•,0_ and (**PBP**)_•,3_ are the products of entries in column **P**_•,3_ and (**BP**)_•,0_ or (**BP**)_•,3_. In other words, the nonzero columns of **PBP** are

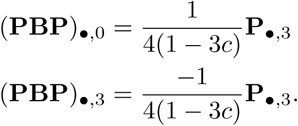

The generating matrix is given by **G=PBP** as in eq. 43.

### The matrix exponential Π(*t*)

To compute the exponential *e*^*t***G**^, we first note that **G**^2^ = −**G***/*[4(1 − 3*c*)]. In general, for *n* > 0,

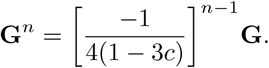

We can then derive the matrix exponential *e*^*t***G**^, converting *t* to units of generations:

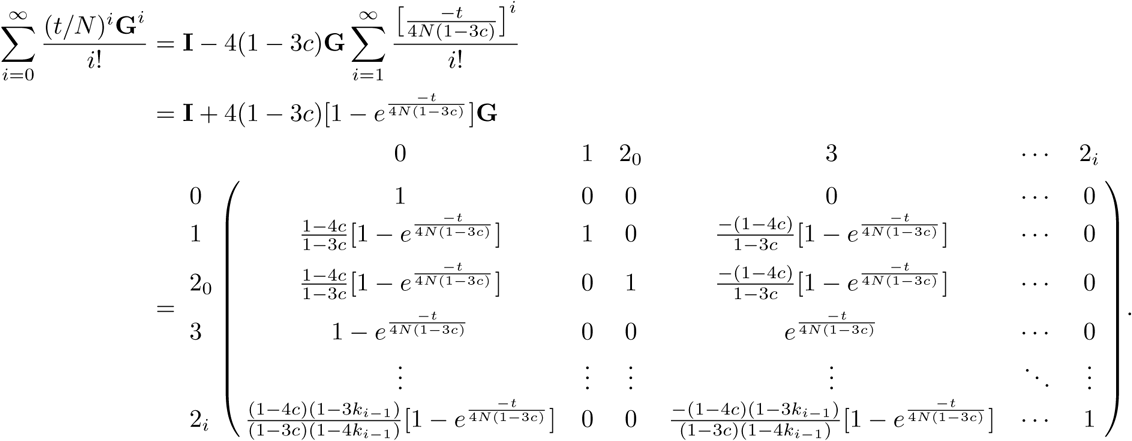

We take the product **P***e*^*t***G**^, using eq. 42 for **P**, to produce Π(*t*), eq. 44.

